# Lack of *Kcnn4* improves mucociliary clearance in muco-obstructive lung disease

**DOI:** 10.1101/2020.05.06.079707

**Authors:** Génesis Vega, Anita Guequén, Amber R. Philp, Ambra Gianotti, Lilian Arzola, Manuel Villalón, Olga Zegarra-Moran, Luis J.V. Galietta, Marcus A. Mall, Carlos A. Flores

**Author notes:** Corresponding author: Carlos A. Flores, Centro de Estudios Científicos (CECs). Valdivia, 511046, Chile.

## Abstract

Airway mucociliary clearance (MCC) is the main mechanism of lung defense keeping the airways free of infection and mucus obstruction. Airways surface liquid (ASL) volume, ciliary beating and mucus are central for proper MCC, and are critically regulated by sodium (Na^+^) absorption and anion secretion. Impaired MCC is a key feature of muco-obstructive disease. The calcium-activated potassium (K^+^)channel KCa.3.1, encoded by the *Kcnn4* gene, participates in intestinal ion secretion and previous studies showed that its activation increase Na^+^ absorption in airway epithelia, suggesting that hyperpolarization induced by KCa3.1 was sufficient to drive Na^+^ absorption. However, its role in airway epithelial function is not fully understood. We therefore aimed to elucidate the role of KCa3.1 in MCC in a genetically engineered mouse model. We show that KCa3.1 inhibition reduced Na^+^ absorption in mouse and human airway epithelium. Furthermore, the genetic deletion of *Kcnn4* enhanced cilia beating frequency (CBF) and MCC *ex vivo* and *in vivo*. *Kcnn4* was silenced in the *Scnn1b*-transgenic mouse (*Scnn1b*^tg/+^), a model of muco-obstructive lung disease triggered by increased epithelial Na^+^-absorption, leading to improvements in MCC and reduction of Na^+^-absorption. KCa3.1 deletion did not change the amount of mucus but did reduce mucus adhesion, neutrophil infiltration and emphysema. Our data support that KCa3.1 inhibition attenuated muco-obstructive disease in the *Scnn1b*^tg/+^ mice. K^+^-channel modulation may be a novel therapeutic strategy to treat muco-obstuctive lung diseases.

## INTRODUCTION

Mucociliary clearance (MCC) is a key process sustaining airway innate defense and it proper function relies on coordinate regulation of epithelial ion and fluid transport, mucus homeostasis and ciliry beating (Mall, 2008; Bustamante-Marin & Ostrowski, 2017). Disruption of any of these epithelial functions reduces airway clearance capacity as observed in asthma, chronic obstructive pulmonary disease (COPD), idiopathic pulmonary fibrosis or primary ciliary dyskinesia, all lung diseases characterized by mucus accumulation, airway obstruction, infections and progressive bronchiectasis (Fahy & Dickey, 2010; Seibold *et al.*, 2011; Tilley *et al.*, 2015). Cystic fibrosis (CF) is, the most common genetic disorder in humans, the “flagship” of all muco-obstructive diseases. Mutations in the *CFTR* gene impede anion transport through the CFTR-protein, which reduce lung function dramatically (Harun *et al.*, 2016). The absence of chloride secretion into the airway lumen dehydrate the airway surface liquid (ASL) impairing cilia movement while absent HCO_3_^−^secretion (that can be secondary to Cl^−^/HCO_3_^−^exchange or directly through CFTR) will affect mucins deployment making it sticky and hard to transport. Reduced HCO_3_^−^secretion also produces acidic pH which in turn decrease activity of ASL anti-microbial molecules, favoring infection settlement (Pezzulo *et al.*, 2012).

Although there is still an ongoing discussion regarding the existence of increased Na^+^ absorption in CF tissues (Donaldson & Boucher, 2003; Chen *et al.*, 2010; Itani *et al.*, 2011; O’Donoghue *et al.*, 2013), its contribution to airway malfunction is tangible in a transgenic animal model with epithelial Na^+^ channel (ENaC) hyper-activity in the airways, that induced a muco-obstructive and inflammatory lung disease phenotype that shares key features with CF and COPD (Mall *et al.*, 2004, 2008). This observation goes in hand with the attempts to use ENaC-channel blockers as a therapy for CF patients that have shown improvements in mucus clearance but of short duration (Hofmann *et al.*, 1997; Hirsh *et al.*, 2004, 2008) or with occurrence of renal side-effects (O’riordan *et al.*, 2014). Genetic silencing using siRNA-based strategies directed against ENaC, produced long-lasting increase in ASL and cilia beating frequency of human bronchial epithelial cells from CF patients (Gianotti *et al.*, 2013; Tagalakis *et al.*, 2018), indicating again that reduction of Na^+^ absorption is of potential benefit in CF and hence other muco-obstructive disorders.

The mechanism for Na^+^-absorption, as described by Koefoed-Johnsen and Ussing in the late 50ties, require basolateral K^+^ channel activity to sustain the electrical membrane gradient (Koefoed‐ Johnsen & Ussing, 1958), so Na^+^ absorption in the airways can hypothetically be regulated by basolateral K^+^ channel function. Tens of K^+^-channels are expressed in the airway epithelium, but in most cases their role in ion transport mechanisms of the lower airway epithelium has not been tested (Bardou *et al.*, 2009).

In this study, we determined the role of the KCa3.1 channel using inhibitors and *Kcnn4* silencing in ion transport and mucociliary clearance (MCC). Using an animal model of CF/COPD-like muco-obstructive lung disease, we generated a double mutant animal to test the hypothesis that *Kcnn4* silencing ameliorates airway disease.

## METHODS

### Reagents

Unless stated otherwise, all reagents were obtained from Sigma-Aldrich, with the exception of TRAM-34 that was bought from TOCRIS.

### Animals

Mice were housed at the CECs animal facility. *Kcnn4*^−/−^animal generation and their genotyping have been described (Begenisich *et al.*, 2004). The *Scnn1b*^tg/+^ mouse was backcrossed to the C57BL/6J background as previously described (Johannesson *et al.*, 2012). Animals were maintained in the Specific Pathogen Free mouse facility of Centro de Estudios Cientificos (CECs) with access to food and water ad libitum. 6-week-old male or female mice were used (C57BL/6J). All experimental procedures were approved by the Centro de Estudios Científicos (CECs) Institutional Animal Care and Use Committee and follow the relevant guidelines and regulations.

### Primary human bronchial epithelial cell culture

The procedures for isolation and culture of human bronchial epithelial cells (HBECs) were described in detail in a previous study (Scudieri *et al.*, 2012). Briefly, HBECs were cultured in flasks in a serum-free medium (LHC9/RPMI 1640). After 4–5 passages, cells were seeded at high density (500,000/cm^2^) on Snapwell 3801 porous inserts (Corning Costar). Twenty-four hours after seeding, the medium was switched to DMEM/F12 (1:1) plus 2% New Zealand fetal bovine serum (Thermo Fisher Scientific), hormones, and supplements. The medium was replaced daily on both sides of the permeable supports for up to 6-7 days (liquid-liquid culture, LLC). Subsequently the apical medium was removed and the cells received nutrients only from the basolateral side (air-liquid culture, ALC). This condition favored further differentiation of the epithelium. Cells were maintained under ALC for 2 weeks before the experiments were performed. The collection of bronchial epithelial cells and their study to investigate the mechanisms of transepithelial ion transport were specifically approved by the Ethics Committee of the Istituto Giannina Gaslini following the guidelines of the Italian Ministry of Health (updated registration number: ANTECER, 042-09/07/2018). Each patient provided informed consent to the study using a form that was also approved by the Ethics Committee.

### Ussing chamber experiments

Tracheae were placed in P2306B of 0.057 cm^2^ tissue holders and placed in Ussing chambers (Physiologic Instruments Inc., San Diego, CA, USA). Tissues were bathed with bicarbonate-buffered solution (pH 7.4) of the following composition (in mM): 120 NaCl, 25 NaHCO_3_, 3.3 KH_2_PO_4_, 0.8 K_2_HPO_4_, 1.2 MgCl_2_, 1.2 CaCl_2_ and 10 D-Glucose, gassed with 5% CO2–95% O2 and kept at 37°C. The transepithelial potential difference referred to the serosal side was measured using a VCC MC2 amplifier (Physiologic Instruments Inc.). The short-circuit currents (I_SC_)were calculated using Ohm’s law as previously described (Gianotti *et al.*, 2016). Differences (Δ I_SC_), were calculated from I_SC_ after minus before the addition of drugs. The experiments were performed in the presence of 10 μM amiloride in the apical side to block Na^+^ absorptive currents. The 1 μM forskolin (FSK) + 100 μM 3-isobutyl-1-methylxanthine (IBMX) cocktail was used to stimulate cAMP-dependent anion secretion, and 100 μM uridine-5′-triphosphate (UTP) to evoke Ca^2+^-activated anion secretion. For experiments on HBECs Snapwell supports carrying differentiated bronchial epithelia were mounted in a vertical chamber resembling an Ussing system with internal fluid circulation. Both apical and basolateral hemichambers were filled with 5 ml of a Krebs bicarbonate solution containing (in mM): 126 NaCl, 0.38 KH2PO4, 2.13 K2HPO4, 1 MgSO4, 1 CaCl2, 24 NaHCO3, and 10 glucose. Both sides were continuously bubbled with a gas mixture containing 5% CO2 – 95% air and the temperature of the solution was kept at 37 °C. The transepithelial voltage was short-circuited with a voltage-clamp (EVC-4000, World Precision Instruments) connected to the apical and basolateral chambers via Ag/AgCl electrodes and agar bridges (1 M KCl in 1% agar). The offset between voltage electrodes and the fluid resistance were canceled before experiments. The short-circuit current was recorded with a PowerLab 4/25 (ADInstruments) analogical to digital converter connected to a computer.

### Cilliary beating frequency

Primary cultures of mouse airway epithelia were analyzed as previously described (Zhao *et al.*, 2012). For all experiments, the central area on each ALI culture was imaged, and the digital image signal was routed from the camera directly into a digital image acquisition board (National Instruments, Austin, TX, USA) within a Dell XPS 710 workstation (Dell, Inc., Round Rock, TX, USA). Images were analyzed with virtual instrumentation software [Sissons-Ammons Video Analysis (SAVA); National Instruments], which is highly customized to quantify CBF. All of the recordings in the present experiments were made at 630x magnification. Whole-field analysis was performed, with each point measured representing one cilium. For each sample, the reported frequencies represent the arithmetic means of these values. Changes in CBF are normalized to the basal rate and reported as a ratio of stimulated/basal.

### Plastic beads clearance and *in-vivo* MCC

Speed of polystyrene beads in trachea samples was determined with some modifications of the previously described method (Ousingsawat *et al.*, 2009). Briefly, mice were deeply anesthetized via intraperitoneal injection of 120 mg/kg ketamine and 16 mg/kg xylazine, exsanguinated, the trachea was isolated and mounted using insect’s needles in an extra thick blot paper (Bio-Rad) which was perfused with Ringer solution (gassed with 95%/5% O_2_/CO_2_) at a rate of 1 ml/min, at 37 °C and maintained in a humidified chamber. Polystyrene black dyed microspheres (diameter 6 μm, 2.6% solid-latex, Polybead, Polyscience Inc., Warrington, PA) were washed and diluted with physiological solution (0,5% latex) and 4 μl of particle solution were added onto the bronchial edge of the trachea. Particle transport was visualized by images every 5s (2-5 minutes sampling) of at least 3 different fields using a Motic camera (Moticam 5.0). Stored images were measured to determine the velocity of single particles using NIH ImageJ software and MCC was expressed in μm/sec. Allergen clearance from the whole lung was evaluated by the elimination of fluorescently-labeled ovoalbumin (alexa fluor 647) after intratracheal instillation as previously described (Mall *et al.*, 2008). 1 mg/ml. Ovalbumin was labelled with Alexa Fluor 647 with the microscale Protein labelling kit (Alexa Fluor 647 Microscale Protein Labelling Kit (Life Technologies, Darmstadt, Germany). Intratracheal instillation of 2.5 μg of OVA dissolved in 20 μl PBS was performed and held over 6 hours. To determine allergen clearance in the whole lung, mice were anesthetized via intraperitoneal injection with 10 mg/kg of ketamine and 16 mg/kg of xylacine. Lung was extracted after 6 hours. To determine the allergen clearance, lung was mixed with PBS and protease Inhibitor 1X (Roche, Basel, Switzerland). The lung was homogenized and fluorescence intensity was read in a microplate reader. Clearance was calculated using the formula:

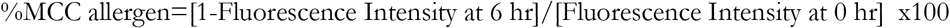

The fluorescence intensity at baseline (t = 0) (arbitraries units) should not interfere between treatments.

### Lung tissue isolation, bronchoalveolar lavage fluid, and histological analysis

Anesthetized mice were euthanized by exsanguination and the chest cavity was opened to ligate the left main bronchus. A blunt needle (20 gauge) was inserted through a small incision in the upper trachea and tied in place with 3.0 silk. After ligation of the left main-stem bronchus, BAL was performed on the right lobes by instilling a volume of sterile PBS (137 mM NaCl, 2.7 mM KCl, 10 mM Na2HPO4, 2 mM KH2PO4) at room temperature determined by the formula:

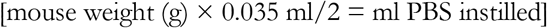

BAL was performed by gently injecting and retrieving the PBS volume three times. This procedure was carried out a second time with an equal volume of PBS and fractions were pooled. Return volume was consistently > 80% of the instilled volume. BAL cells were pelleted by centrifugation at 300 × g for 5 min at 4°C and the cell-free supernatant (BAL fluid or BALF) was collected and stored at −80°C with protease inhibitors. BAL cells were resuspended in 50 μl PBS and 10 μl were counted with a hemocytometer (diluted in 10 μl Tripan blue). 40 μl were diluted in 160 μl and displaced in Cytospin slides (StatSpin Cytofuge 2, Norwood, MA), air dried, and stained with modified Giemsa for differential cell counts of at least 200 cells per slide. After BAL, the left and right lung were immersed in 10% neutral-buffered formalin to prevent the dislodging of airway luminal contents. The right lung was used to evaluate the mean linear intercept and the left lung to quantify mucus density and attachment using Image J.

### Lung histology and mucus morphometry

Morphometric analyses of airway mucus obstruction were performed in non-inflated, immersion fixed left lungs. Lungs were removed through a median sternotomy, fixed in 4% buffered formalin, and embedded in paraffin. Left lung was sectioned transversally at the level of the proximal intra-pulmonary main axial airway near the hilus (proximal airways), and at the distal intra-pulmonary axial airway, at 1500 μm distal to the hilus (distal airways). To quantify secreted mucus, morphometric analysis of stained sections was carried out by determining mucus volume density using CellF Software, as previously described (Mall *et al.*, 2008). Briefly, the length of the basal membrane of the airway epithelium was measured by the interactive image measurement tool, and the AB-PAS positive surface area within this boundary was measured by phase analysis according to the automatic threshold settings of the software. The volume density of intraepithelial mucus, representing the volume of lumen mucus content per surface area of the mucus basal membrane (nl/mm^2^), was determined from the intraepithelial surface area of Alcian Blue-Periodic Acid Schiff staining (AB-PAS) positive mucus and the length of the basal membrane of the airway epithelium. Luminal mucus content relative to the luminal area was determined in a similar fashion.

### mRNA isolation and cDNA synthesis from epithelial cells

Mice were killed by cervical dislocation and tracheal tissues were immediately extracted, tracheas were incubated with Pronase 30 μM at 37°C during 30 minutes. Trachea was placed in DMEM 10 mM D-Glucose and epithelium was isolated by scraping with tweezers and further homogenized in 250 μl of Trizol (TRIzolTM Reagent) and RNA isolated following the manufacturer’s instructions. The dried pellet of RNA was re-suspended with 35 μl of nuclease free water, and stored at –80°C. DNA contamination was avoided using DNase treatment. The concentration and integrity of the RNA were determined by spectrophotometry. Total RNA was reverse transcribed into cDNA using the Superscript III RTPCR System according to the manufacturer’s recommendations. cDNA synthesis was performed on 2 μg RNA. cDNA integrity was checked using specific primers to cyclophilin. 50 ng template cDNA was added to the reaction mixture. *Cyclophilin* (*Ppia*) amplification was performed starting with a 5 min template denaturation step at 95°C, followed by 30 cycles of denaturation at 95°C for 30 s and combined primer annealing/extension at 55°C. The relative brightness intensity of ethidium bromide-stained bands resolved on a 1.5% agarose gel was evaluated.

### Real Time PCR

Quantification of *Scnn1a*, *Scnn1b*, *Scnn1g*, and *Prss8* expression was performed using SYBR Green detection in a LightCycler PCR machine according to the manufacturer’s instructions. We determined the PCR efficiency of each individual assay by serially measuring 100 ng cDNA from a pool of epithelia in triplicate. Only CT values < 40 were used for calculation of the PCR efficiency. All PCRs displayed an efficiency between 96 and 100%. Amplifications were performed starting with a 3 min template denaturation step at 94°C, followed by 45 cycles of denaturation at 94°C for 20 s and combined primer annealing/extension at the gene specific primer temperature for 30 s. All samples were amplified in triplicate and the mean was obtained for further calculations. Relative-fold changes in target gene expression were quantified by the previously reported ΔΔCT method (Livak & Schmittgen, 2001). Briefly, CT values were obtained for individual samples using the Rotor-Gene 6000 software 1.7 (Corbett Life Science Pty Ltd., Sydney, Australia), where the targets and reference (Cyclophilin) had the same cDNA concentration. ΔCT was calculated by subtracting the CT (target – reference). Primers and annealing temperatures are given in Table 1.

**TABLE 1.**
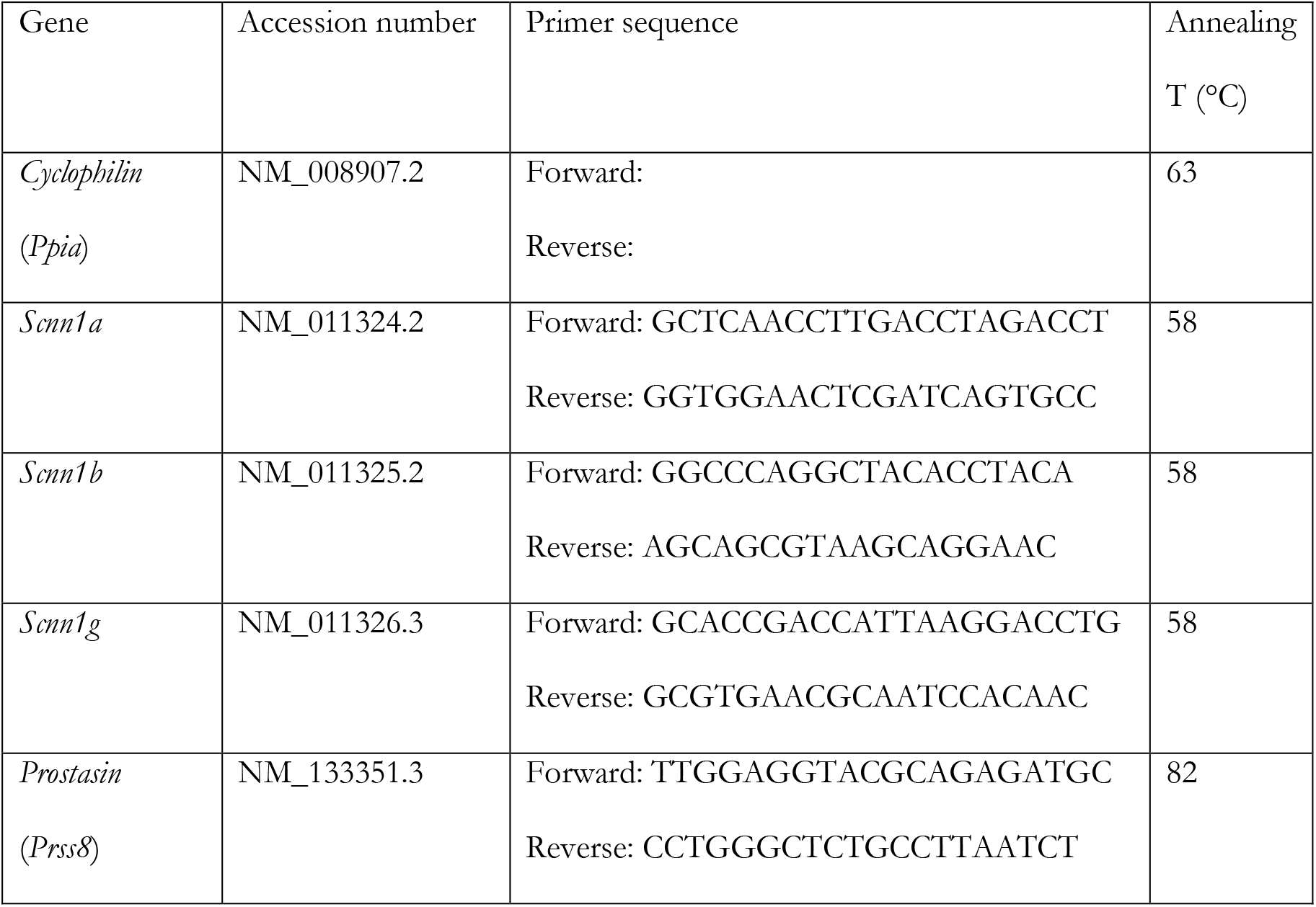
Primer sequences and annealing temperatures.

### Statistical analysis

All statistical analyses were performed using the SigmaPlot V12.1. Tests are annotated in figure and table legends.

## RESULTS

### Na+-absorption and calcium-activated anion secretion is dependent on KCa3.1 activity in mouse and human airway epithelium

First we aimed to determine if the genetic deletion of *Kcnn4* affected electrogenic ion transport in the airway epithelium of mouse trachea. Representative traces of freshly excised wild type and *Kcnn4*^−/−^tracheal tissues are presented in Figs 1A and 1B. We observed that the transepithelial potential (V_te_) was significantly reduced in the *Kcnn4*^−/−^trachea when compared with wild type (Fig 1C). We observed no changes in R_te_ (Fig 1D) or basal I_sc_ (Fig 1E). The amiloride-sensitive current was significantly reduced in the *Kcnn4*^−/−^tissues (Fig 1F). Both the amiloride-insensitive (Fig 1G) and cAMP-induced anion secretion (Fig 1H) remained unaffected by *Kcnn4* silencing. The peak response of Ca^2+^-activated anion secretion induced by UTP (Fig 1I) and the plateau phase (5 minutes after UTP addition; Fig 1J) were reduced. Bio-electrical parameters are summarized in Table 2. We also tested if the silencing of another K^+^ channel that is also important for anion secretion, the KCNQ1/KCNE3, affected Na^+^ absorption using *Kcne3*^−/−^tracheas but observed no effect (Table 2).

**TABLE 2.** Bioelectrical parameters of mouse tracheas.

|  | Wild type (9) | *Kcnn4*<sup>-/-</sup> (9) | *Kcne3*<sup>-/-</sup> (5) | *Scnn1b*<sup>tg/+</sup> (6) | Double mutant (4) |
| --- | --- | --- | --- | --- | --- |
| V<sub>te</sub> (mV) | -3.76 ± 0.8 | -2.04 ± 0.4* | -1.07 ± 0.3* | -3.51 ± 0.6 | -2.24 ± 0.4 |
| R<sub>te</sub> (Ω cm<sup>-2</sup>) | 91 ± 12 | 75 ± 6 | 64 ± 7 | 59 ± 7 | 53 ± 2 |
| I<sub>sc</sub> (μA cm<sup>-2</sup>) | -39.4 ± 5 | -28.1 ± 5 | -14 ± 4* | -65 ± 3* | -53.1 ± 9 |
| Amiloride sensitive<br>(μA cm<sup>-2</sup>) | -17.6 ± 3.8 | -6.2 ± 1.8* | -11.5 ± 1.4 | -25.2 ± 1.7* | -12.7 ± 3§ |
| Amiloride Insensitive<br>(μA cm<sup>-2</sup>) | -21.8 ± 3.8 | -23.0 ± 4.6 | -2.7 ± 2.6* | -41.8 ± 3.6* | -40.3 ± 6.7* |
| IBMX+FSK induced<br>(μA cm<sup>-2</sup>) | -12.8 ± 1.9 | -9.7 ± 1.4 | n.d. | n.d. | n.d. |
| UTP induced peak<br>(μA cm<sup>-2</sup>) | -91.1 ± 12 | -58.2 ± 9.6* | n.d. | n.d. | n.d. |
| UTP induced plateau<br>(μA cm<sup>-2</sup>) | -26.8 ± 4.7 | -10.1 ± 4.1* | n.d. | n.d. | n.d. |

\* Indicates p<0.05 vs Wild type, Rank sum test. Other comparisons can be found on Figures 1 and 3. § Indicates p<0.05 vs *Scnn1b*<sup>tg/+</sup>, Rank sum test.

**FIGURE 1.**
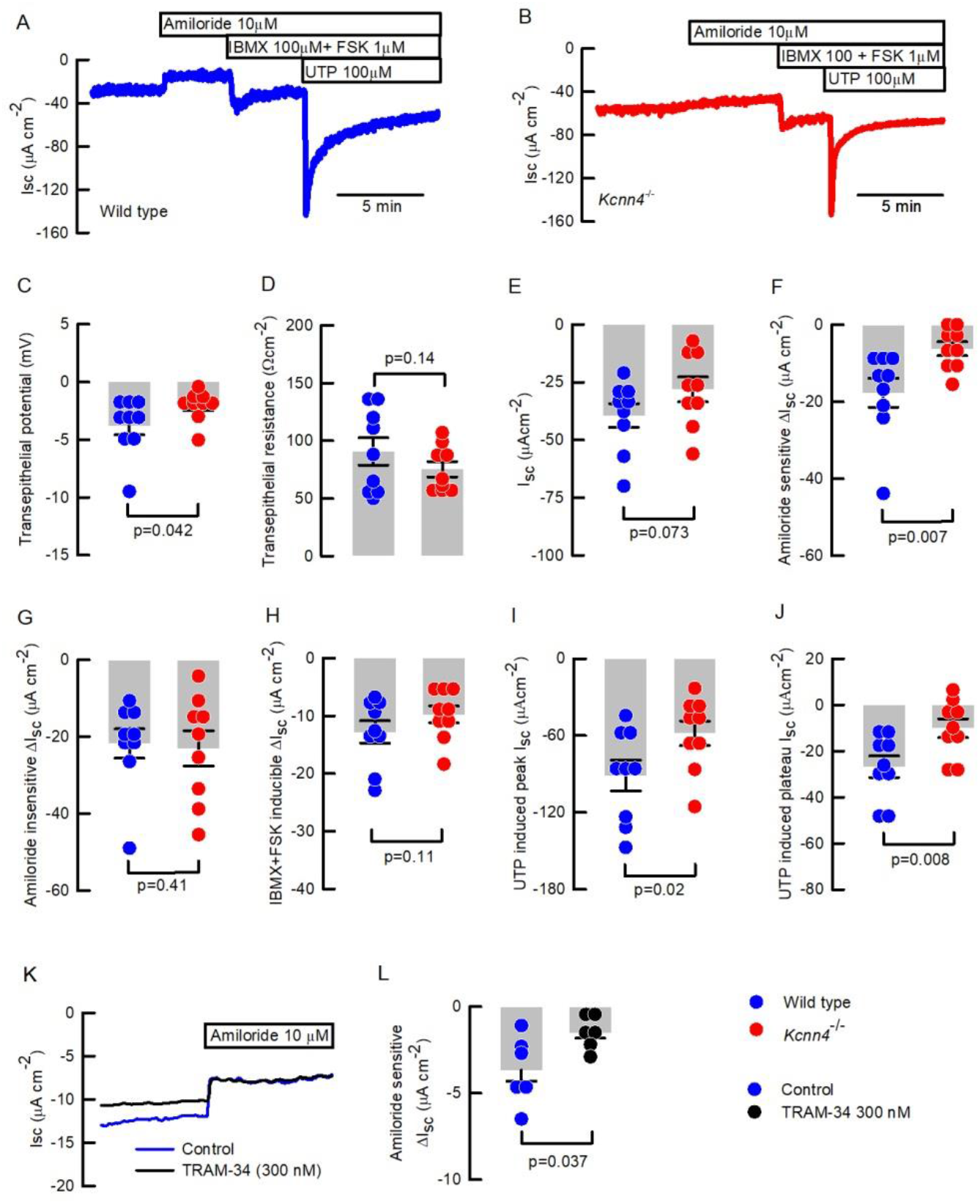
KCa3.1 participates in sodium absorption and anion secretion of mouse and human epithelium. Representative short-circuit current traces of (A) Wild type and (B) *Kcnn4*^−/−^mouse tracheas used to calculate the following: (C) V_te_, (D) R_te_, (E) I_SC_, (F) Amiloride sensitive current, (G) Amiloride-insensitive current, (H) cAMP-induced anion current and Ca^+2^-activated anion current at (I) peak or (J) plateau phases. (K) Representative Ussing chamber recordings of HBECs incubated with TRAM-34 and controls showing amiloride addition. (L) Summary of TRAM-34 effect on Amiloride-sensitive sodium absorption. Statistical differences were calculated using Rank Sum test. Detailed values including data for *Kcne3*^−/−^are included in Table 2.

To determine if KCa3.1 inhibition mimics the observations in mouse trachea we next tested known inhibitors of KCa3.1 channels clotrimazol and TRAM-34 in HBEC monolayers mounted in Ussing chambers. Clotrimazol (30μM) did not affect on amiloride-sensitive Na^+^ absorption but produced a significant reduction in both cAMP and Ca^2+^-activated anion secretion (Sup Fig 1A-C). TRAM-34 produced a significant reduction of the amiloride-sensitive current when used at 300 nM (Fig 1K and L), with no effect on cAMP or Ca^2+^-activated anion secretion (Sup Fig 1B & C). When TRAM-34 was used at 100 nM there was no effect on any of the measured parameters (Sup Fig 1A-C).

To discard that the reduction in the amiloride-sensitive current is produced by a reduced expression of ENaC sub-units or by a reduction in the epithelial protease and ENaC activator prostasin (*Prss8*), we performed qRT-PCR analysis and demonstrated that all ENaC subunits and *Prss8* gene expression were unaltered after *Kcnn4* silencing (Sup Fig 2).

**FIGURE 2.**
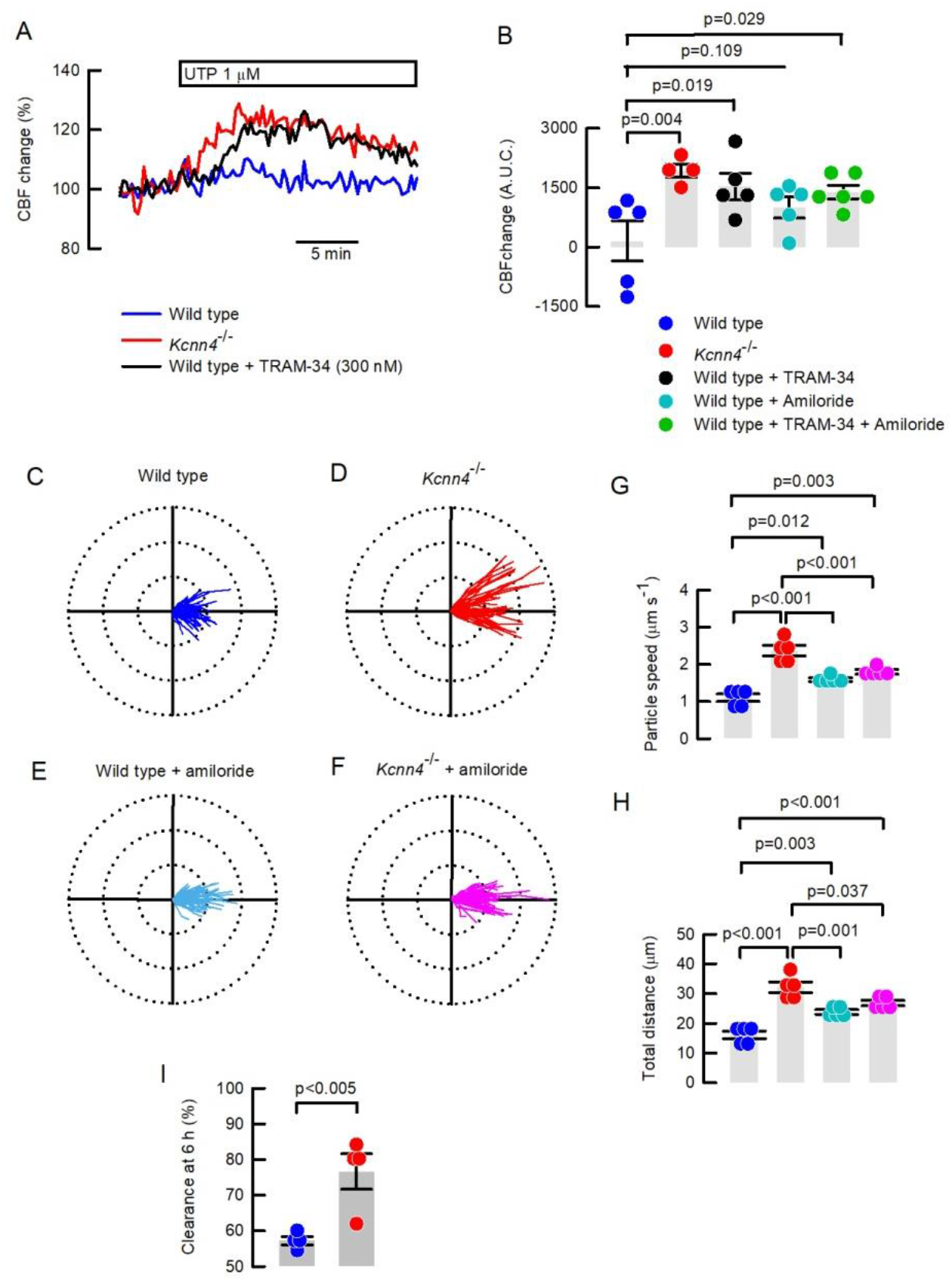
Mucociliary clearance is enhanced after KCa3.1 inhibition in the mouse. (A) CBF changes in *Kcnn4*^−/−^or wild type cells treated with TRAM-34 (100 nM) after UTP stimulation of tracheal epithelial cell explants. (B) Quantification of the area under the curve (A.U.C.) of CBF recordings including wild type explants treated with amiloride (10μM) and/or TRAM-34. Polar plots (75 μm radius) of the trajectory of beads placed in (C) wild type (80 beads), (D) *Kcnn4*^−/−^(68 beads), (E) wild type + 10 μM amiloride (84 beads) and (F) *Kcnn4*^−/−^+ 10 μM amiloride (81 beads) tracheas, from 5 different experiments each. Calculated (G) speed of particles and (H) total distance from the experiments shown in C-F. Differences were calculated using ANOVA on ranks. (I) *in vivo* lung clearance of fluorescently-labelled OVA is increased in the *Kcnn4*^−/−^mouse. The difference was calculated using Rank Sum test.

### KCa3.1 inhibition increases ciliary beating frequency and clearance in mouse airways

To test if reduced Na^+^ absorption affected mucociliary clearance we studied changes in UTP-induced ciliary beating frequency (CBF) determined in *Kcnn4*^−/−^or wild type mouse airway cells cultures incubated with TRAM-34 or/and amiloride. As observed in Fig 2A genetic silencing or TRAM-34 increased UTP-induced CBF. The summary in Fig 2B also includes experiments with amiloride with or without addition of TRAM-34. Amiloride alone did not significantly increase CBF, and only achieved this in combination with TRAM-34.

Since CBF experiments must be performed in submerged conditions, we tested the clearance of plastic beads in freshly excised mouse tracheas whose mucosal side was exposed to air in a humidified chamber. Figure 2C and D show polar plots for wild type and *Kcnn4*^−/−^tissues where an increase in the distance covered by the beads is clearly seen. Detailed analysis of the movement of the beads showed that *Kcnn4* silencing induced a significant increase in the average speed (Fig 2G) and also in the total covered distance (Fig 2H). To evaluate if inhibition of Na^+^ absorption affected the speed and distance travelled by the plastic beads we used amiloride and observed an increase in both parameters in wild type tracheas (Fig 2G and H), as previously demonstrated (Joo *et al.*, 2016), nevertheless, when amiloride was tested in the *Kcnn4*^−/−^tracheas a decrease in MCC parameters was observed compared to non-treated *Kcnn4*^−/−^tracheas, but still maintained higher MCC parameters than control wild type tissues (Fig 2G and H). Finally, we tested whole lung clearance *in vivo*, measuring the clearance of fluorescent-labeled OVA and found a higher clearance percentage in the *Kcnn4*^−/−^animals correlating with the *ex vivo* observations and demonstrating that *Kcnn4* silencing increases MCC (Fig 2I).

### Genetic silencing of *Kcnn4* reduces Na^+^-absorption and increases MCC in the *Scnn1b*^tg/+^ mouse

To test if the improved MCC observed in the *Kcnn4*^−/−^mouse might be of potential benefit, we mated the *Kcnn4*^−/−^animals with the *Scnn1b*^tg/+^ mouse that is affected by severe inflammatory and muco-obstructive airway disease. First, we studied if *Kcnn4* silencing induced changes in epithelial Na^+^-absorption and airway clearance. Ussing chamber experiments demonstrated that amiloride-sensitive Na^+^ absorption was significantly decreased in double mutants compared to tracheas from the *Scnn1b*^tg/+^ mouse (Fig 3A and B), reaching values similar to those observed in the wild types (Fig 3A, B and Table 2). The clearance of beads also showed increased values for speed and distances (Fig 3C-F), indicating that *Kcnn4* silencing improved MCC in the muco-obstructive lung disease model.

**FIGURE 3.**
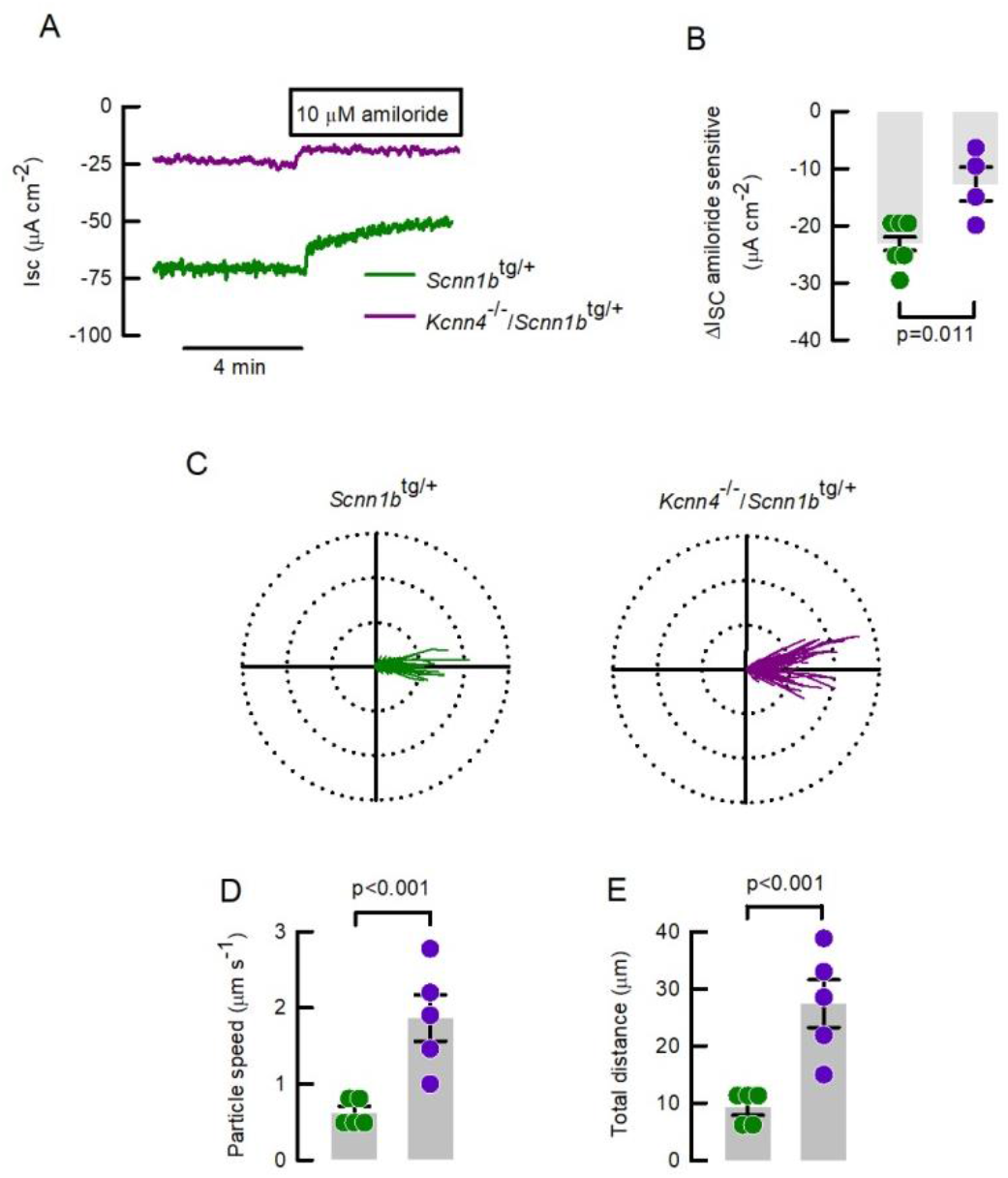
The genetic silencing of *Kcnn4* reduces sodium absorption and increases MCC in the *Scnn1b*^tg/+^ mouse trachea. (A) Representative I_SC_ traces showing the extent of amiloride-sensitive currents in the *Scnn1b*^tg/+^ and double mutant mice trachea. (B) Summary of amiloride-sensitive currents as shown in (A), p value calculated using ANOVA on ranks. Polar plots (75 μm radius) of the trajectory of beads placed in (C) *Scnn1b*^tg/+^ (222 beads) and double mutants (126 beads), from 5 different experiments each. Calculated (D) speed of particles and (E) total distance from the experiments shown in C-F. Differences were calculated using ANOVA on ranks.

### Genetic silencing of *Kcnn4* prevents airway inflammation and lung damage in the *Scnn1b*^tg/+^ mouse

After *Kcnn4* silencing lung edema in the *Scnn1b*^tg/+^ mouse was reduced to levels similar to the control animals (Fig 4A) and concomitantly, we observed less damage associated with emphysema (Fig 4B and C). Analysis of BALF was performed to quantify the absolute numbers and relative distribution of immune cell types present in the airways of the animals. As previously observed (Mall *et al.*, 2004), the *Scnn1b*^tg/+^ mouse showed increased amounts of total cells (Fig 4D), corresponding to macrophages (Fig 4E) and neutrophils (Fig 4F). The silencing of *Kcnn4* did not affect the number of total cells or macrophages but reduced the number of neutrophils in the *Scnn1b*^tg/+^ airways, an observation that was confirmed with staining and quantification of neutrophils in mucus plugs with LY6G/LY6C (Fig 4G and H). The silencing of *Kcnn4* alone, did not produce changes in the measured parameters (Fig 4A-C, F-H).

**FIGURE 4.**
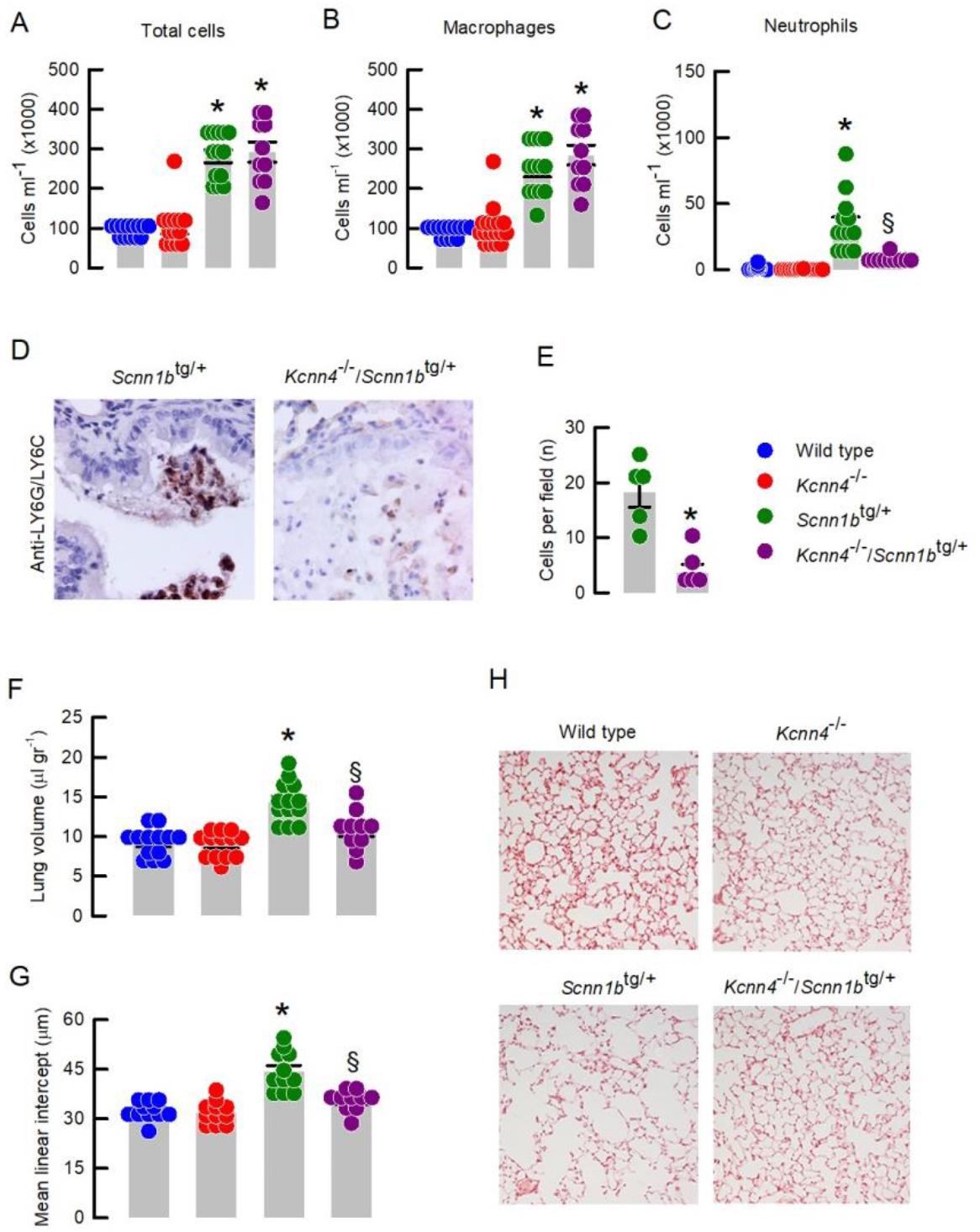
Genetic silencing of *Kcnn4* reduces lung inflammatory disease in mice with muco-obstructive lung disease. (A) Total cells and (B) macrophages quantification in BALF. * indicates p<0.05 vs wild type and *Kcnn4*^−/−^. (C) Neutrophils quantification in BALF. * indicates p<0.05 vs all other groups and § indicates the difference vs *Scnn1b*^tg/+^; ANOVA on ranks. (D) Representative images of LY6G/LY6C immunostaining in mucus plugs. (E) Quantification of LY6G/LY6C positive cells in the *Scnn1b*^tg/+^ and double mutants. * indicates p<0.05 calculated by Rank Sum test. (F) Lung volume and (G and H) emphysema are reduced in in the *Scnnb1*^tg/+^ mouse after the silencing of *Kcnn4*. * indicates p<0.05 vs all other groups and § indicates the difference vs *Scnn1b*^tg/+^; ANOVA on ranks.

To test if the improvement in lung disease correlates with a decrease in mucostasis, we analyzed lung histological samples corresponding to proximal bronchi and distal bronchi stained with PAS. Intraluminal and intraepithelial mucus volume were increased in the *Scnn1b*^tg/+^ mouse compared to wild types and *Kcnn4*^−/−^in both, proximal and distal airways (Figs 5A-B). After *Kcnn4* silencing values were not significantly different from those observed in the *Scnn1b*^tg/+^ mouse, but we observed a trend toward reduced intraluminal mucus obstruction in the distal airways of double mutants *Kcnn4*^−/−^/*Scnn1b*^tg/+^ mice that was not statistically significant based on the number of mice available for these studies (Figs 5A). A closer examination of the tissues lead us to the observation that while mucus in the *Scnn1b*^tg/+^ mouse mostly adhered to the epithelial wall this was not the case in the double mutant airways (Fig 5C). Quantification of the epithelial airway surface in contact with mucus demonstrated that there is a gradient of mucus adhesion to the surface of the epithelium (proximal to distal airway gradient) in the wild type animals (17.4 ± 3% and 0.4 ± 0.1% of epithelial surface attached to mucus; p<0.001) that is maintained in the *Kcnn4*^−/−^tissues (14.6 ± 4.8% and 2.3 ± 1.2%; p=0.02; Fig 5D). Analysis of the *Scnn1b*^tg/+^ tissues showed a complete loss of the proximal to distal airway gradient, as both values were similar and higher in both proximal and distal airways (45.0 ± 6% and 45.1 ± 9.1%; p>0.4). Finally, a recovery of the proximal to distal airway gradient was observed after *Kcnn4* silencing (59.7 ± 10.8% and 24.7 ± 7.4%; p=0.014).

**FIGURE 5.**
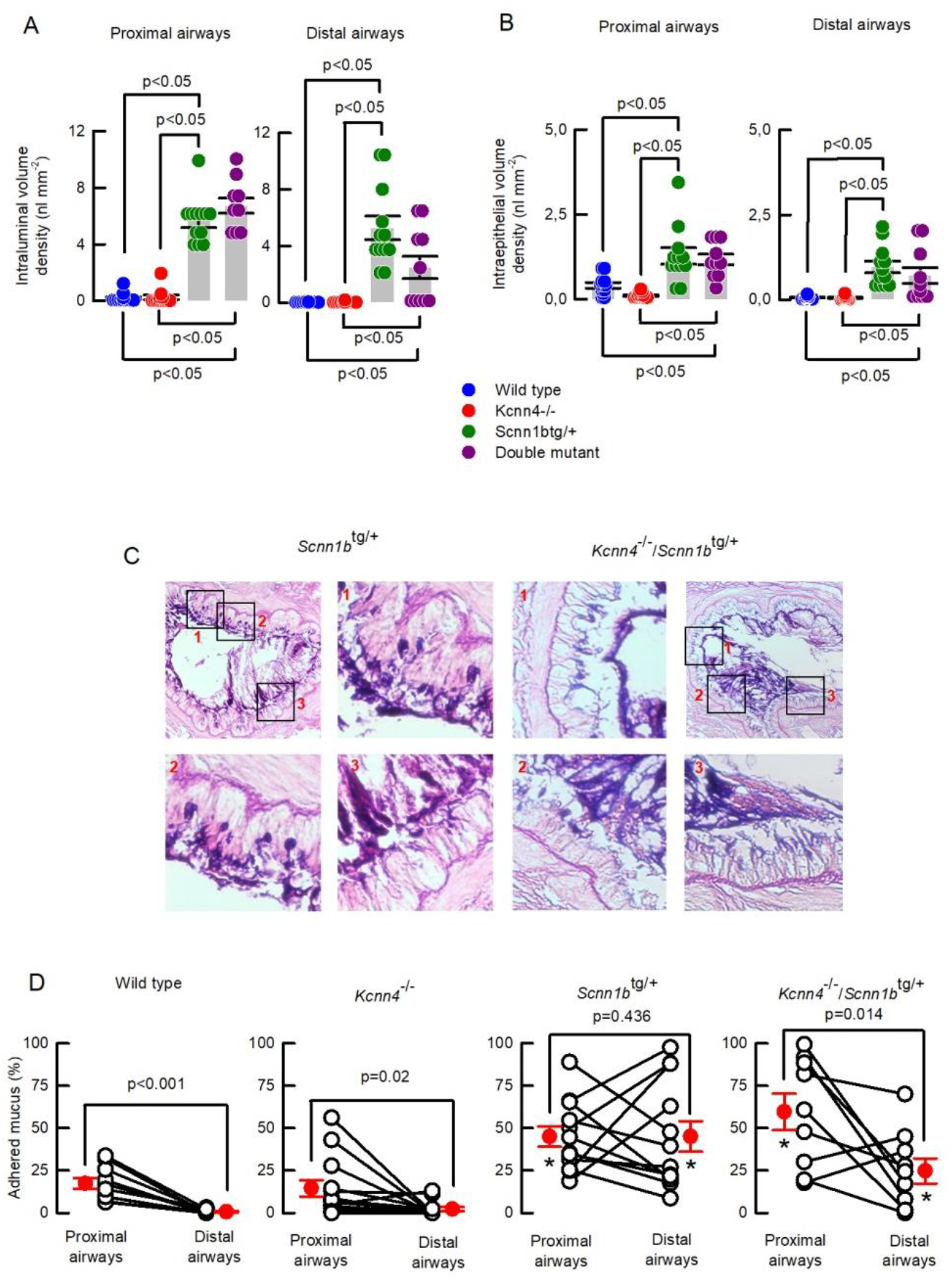
Genetic silencing of *Kcnn4* improved mucus clearance in mouse airways. (A) intraluminal and (B) intracellular mucus volume was determined in the proximal and distal airways. ANOVA on ranks. (C) Representative images of mucus attachment to the epithelial surface for *Scnn1b*^tg/+^ and double mutants. (D) Summary of the percentage of epithelium surface covered my mucus in proximal and distal airways. Only paired samples from the same animal were included. * indicate p<0.05 vs wild type and *Kcnn4*^−/−^ANOVA on ranks. The p values for each proximal vs distal airways comparison were calculated by Rank Sum test.

## Discussion

We identified KCa3.1 as a K^+^ channel responsible for energizing the cell electrical gradient for Na^+^-absorption in human and mouse airway epithelial cells. As predicted from previous observations, reduced Na^+^-absorption might enhance MCC function in the airways of the *Kcnn4*^−/−^mouse, therefore we tested the potential of *Kcnn4* silencing in the amelioration of airway muco-obsturcitve disease in the *Scnn1b*^tg/+^ mouse model. *Kcnn4* silencing successfully decreased inflammatory lung damage, neutrophilic inflammation and improved MCC in the *Scnn1b*^tg/+^ animals. We found that *Kcnn4* silencing also affected anion secretion, however, no signs of muco-obstructive disease were found in this model.

Early observations in patients affected with pseudohypoaldosteronism (due to ENaC inactivating mutations) indicated that ENaC activity was inversely related to ASL airway volume and MCC efficiency (Kerem *et al.*, 1999), suggesting that ENaC might be a druggable target to manage muco-obstructive lung disease. Some rate of success was obtained using ENaC blockers like amiloride and derivatives, but short duration or side effects have hampered further advances in the development of therapies so, new approaches are necessary (Tildy & Rogers, 2015). As described in the 50ties, the Na^+^-absorption mechanism relies on the activity of basolateral K^+^-channels (Koefoed‐Johnsen & Ussing, 1958). Acting in concert to the Na^+^-pump, highly selective basolateral channels allow the extrusion of K^+^ that otherwise retained depolarize the membrane potential, hampering Na^+^-absorption. This is the case in the kidney and intestine where inhibition of basolateral K^+^ channels reduced Na^+^-absorption and Na^+^-coupled metabolite uptake (Turnheim *et al.*, 2002; Hebert *et al.*, 2005; Vallon *et al.*, 2005), further strengthening the role of K^+^ channels on epithelial Na^+^ homeostasis. We envisioned that Na^+^-absorption in the airways can be controlled through basolateral K^+^ channels, but their role in electrolyte transport has not been profusely studied in this tissue (Bardou *et al.*, 2009). With the exceptions of Kir7.1, KCNE3 and KCa3.1 no other K^+^ channels or sub-units have been located in the basolateral membrane of lower airway epithelium (Thompson-Vest *et al.*, 2006; Preston *et al.*, 2010; Villanueva *et al.*, 2015). Experiments with ion channel-inhibitors demonstrated that KCNQ1/KCNE3, using chromanol-293B, and KCa3.1, using clotrimazole, participated in the mechanism of anion secretion of the airways, and experiments in the *Kcne3*^−/−^mouse confirmed its role but no effect on Na^+^-absorption was observed (Mall *et al.*, 2000, 2003; Preston *et al.*, 2010). Our results confirmed these observations, as when using clotrimazole a decrease in anion secretion was observed, but there was no effect in Na^+^-absorption in human cells, similar to what we observed in tracheas from the *Kcne3*^−/−^animals. When using the KCa3.1 opener 1-EBIO we observed an increase in Na^+^-absorption in HBECs supporting a role for KCa3.1 as a regulator of Na^+^-absorption in these cells (Devor *et al.*, 2000), a result comparable to our observations using TRAM-34 in HBECs and tracheas from the *Kcnn4*^−/−^mouse where decreased Na^+^-absorption was recorded. The *Kcnj13*^−/−^mouse bears a lethal phenotype at perinatal age and is affected by severe tracheomalacia, but it is not known if Kir7.1 participates in the anion secretion mechanism and if it relates to the observed phenotype (Yin *et al.*, 2018).

Epithelial anion secretion in the airways is also affected after K^+^-channel silencing that *a priori*, and in the scenario of a muco-obstructive disease, it is unwanted. Both, KCa3.1 and KCNQ1/KCNE3 inhibition produced a significant reduction of Ca^2+^ and cAMP-induced anion secretion respectively (Mall *et al.*, 2000, 2003), but no muco-obstructive phenotype was observed in the *Kcne3*^−/−^(Preston *et al.*, 2010) or *Kcnn4*^−/−^airways as observed in our results. Such reductions in anion secretion are also innocuous in the intestine. For example, *Kcnn4*, *Kcnq1*, *Kcne3* or *Kcnk5* silencing did not produced obstructions or CF-like phenotypes in the intestine of the animals, as expected from genes coding for K^+^-channels with tested roles in epithelial anion secretion (Vallon *et al.*, 2005; Flores *et al.*, 2007; Preston *et al.*, 2010; Julio-Kalajzić *et al.*, 2018). Even though, we don’t have a straight answer for the lack of the muco-obstructive phenotype when *Kcnn4* is silenced, is important to consider that the inhibition of anion secretion in this model is not complete, KCNQ1/KCNE3 activity and basolateral electroneutral-anion influx can still support anion secretion at a sufficient level to maintain MCC function.

But is inhibiting KCa3.1 better than inhibiting ENaC?. Besides the already discussed use of ENaC blockers and other inactivating strategies, animal models have demonstrated that silencing of genes coding for ENaC subunits are lethal, and particularly *Scnn1a* silencing produced lethality due to defective liquid clearance from the neonatal lung (Hummler *et al.*, 1996; Hummler & Vallon, 2005). Nevertheless, we agree with the idea that the key seems to be the dosage of ENaC inhibition, as the heterozygous mutant mice for *Scnn1a* do not show lethality, and residual ENaC activity in *Scnn1b*^−/−^and *Scnn1g*^−/−^is sufficient to overcome the neonatal lung phenotype (Hummler *et al.*, 1996; Barker *et al.*, 1998; Mcdonald *et al.*, 1999). Our results show that after *Kcnn4* silencing ~35% of amiloride-sensitive Na^+^-absorption is still active in mouse trachea, suggesting that the remaining current is sufficient to maintain proper MCC function.

Analysis of the genetic silencing of *Kcnn4* in the *Scnn1b*^tg/+^ animal demonstrated reduced infiltration of neutrophils and decreased emphysemic damage of the respiratory epithelium. We also observed that a gradient of adhered mucus to the epithelium normally occurs, as mucus is covering a larger surface in the proximal bronchi than in the terminal. This gradient was disrupted in the *Scnn1b*^tg/+^ tissues, where similar values for mucus attached to the epithelial surface are observed in proximal and distal bronchi. Even though, we observed a tendency for mucus reduction in the distal airways of double mutants, it was accompanied by a recovery in the gradient of mucus adherence to the epithelial surface, supporting the idea that the mucus hyper-secretory state *per se* is not enough to provoke muco-obstructive disease. For, example animals that overexpress MUC5AC or MUC5B in the lungs do not produce airway obstructions or inflammation, even when in some cases the amount of mucus-protein is almost 20-times higher than in control animals, thus reaffirming the idea that under normal fluid transport the amount of mucus is not an issue (Ehre *et al.*, 2012; Roy *et al.*, 2014). Mucus oversecretion must then be accompanied by airway dehydration and inflammatory signalling to complete the muco-obstructive pathology, and therefore, we conclude that the observed reduction of inflammation and the increased MCC after *Kcnn4* silencing prevents the muco-obstuctive pathology in the double mutant animals irrespective of mucus quantity. Even when mucus hyper-secretion is at the center stage in airway muco-obstructive diseases, animal models have demonstrated that impeding mucin expression is detrimental for animal well-being since mucus is fundamental for lung health (Roy *et al.*, 2014). Studying the same *Scnn1b*^tg/+^ mouse model with additional silencing of *Muc5ac* or *Muc5b*, increased inflammation or no improvements at all in airway disease were found (Livraghi-Butrico *et al.*, 2017). Our results agree with the idea that complete mucus reduction is not recommendable, but interventions that boost MCC, improve mucus quality and/or reduce adhesion of mucus to the epithelium are in the road to limit the seriousness of the disease.

KCa3.1 possesses a high affinity for Ca^2+^ with EC50 values of 95 to 160 nM for the human and mouse channel respectively (Joiner *et al.*, 1997; Vandorpe *et al.*, 1998). Free calcium in the cytoplasm of airway epithelial cells is typically 100 nM (Welsh & McCann, 1985; Lee & Foskett, 2014), but ciliated cells bore a higher concentration at baseline than Club cells (De Proost *et al.*, 2008). Another important functional consideration is that ciliated cells responded increasing Ca^2+^ from 100 nM up to 400 nM when stimulated with a low ATP concentration (0,1-1 μM) (Zhang & Sanderson, 2003), close to what is normally found in the sputum and ASL from humans donors (Donaldson *et al.*, 2000; Li *et al.*, 2006; Anderson *et al.*, 2015). All these evidences argue in favor of the idea that KCa3.1 channels are active during normal conditions in certain cells of the airway epithelium, but further immunolocalization studies are necessary to unveil the exact cell distribution, subcellular localization and protein interactions of KCa3.1 channels in the airways.

Even when decreased Na^+^-absorption after *Kcnn4* silencing fits with expected improvements in MCC, we have to address the fact that some of our results indicate that KCa3.1 functions go beyond epithelial fluid secretion and absorption. Increased CBF after KCa3.1 blockade occurs in submerged conditions, discarding a direct link to increased ASL. The hyperpolarization produced by the opening of KATP channels in the apical membrane increases the speed CBF (Ohba *et al.*, 2013), in a similar fashion, KCa3.1 inhibition might have a hyperpolarizing effect in the apical membrane (by decreasing Na^+^-absorption) boosting CBF. Our results also indicate that KCa3.1 inhibition had a greater effect on CBF than ENaC inhibition. It might be possible that the reduction of anion secretion also favour apical membrane hyperpolarization, a hypothesis that is supported by the more positive Vte recorded in the *Kcnn4*^−/−^tissue that might be due to Na^+^ accumulation in the mucosa and reduced anion secretion. Besides that, KCa3.1 inhibition has been widely proven by different groups to prevent or ameliorate several inflammatory events including those occurring in the airways (Girodet *et al.*, 2013; Lin *et al.*, 2015; Yu *et al.*, 2017). Since our model is a systemic null-animal other might also participate in the reduction of inflammatory damage after *Kcnn4* silencing. Such is the case of neutrophils whose number is reduced in the *Kcnn4*^−/−^lungs affected by acute lung injury (Henriquez *et al.*, 2016). Similar results were obtained in the CF *Cftr*^ΔF508/ΔF508^ mouse, which after *Kcnn4* silencing almost completely reduced lethality associated with intestinal obstructions, but was not accompanied by improvements in intestinal anion secretion (Philp *et al.*, 2018). Finally, the use of KCa3.1 inhibitors must be taken with caution as off-targets are reported (Schilling & Eder, 2007; Agarwal *et al.*, 2013; Bonito *et al.*, 2016), and in our case, the use of two different inhibitors (clotrimazol and TRAM-34) resulted in non-comparable results in Ussing chamber experiments.

Even though patients affected by CF have found therapies in molecules that help correct mutant-CFTR malfunctions (Heijerman *et al.*, 2019; Middleton *et al.*, 2019), and new such molecules are being discovered (Pedemonte *et al.*, 2020), almost 10% of CF patients are still left with no such treatment. The potential use of KCa3.1 inhibitors correspond to what has been called a “mutation-agnostic” treatment for CF, but also seems suitable for easing other muco-obstructive diseases affecting humans. The absence of severe phenotypes in the *Kcnn4*-null animal (Begenisich *et al.*, 2004; Si *et al.*, 2006; Flores *et al.*, 2007; Grgic *et al.*, 2009) suggest the use of KCa3.1 inhibitors may be safe in human diseases.

## Acknowledgements

We want to express our gratitude to the CECs-animal facility personnel for their untiring work. Supported by FONDECYT 1151142 (C.A.F.). CECs is funded by the Centers of Excellence Base Financing Program of CONICYT. M.A.M. has been supported by grants from the German Federal Ministry of Education and Research (82DZL004A1), the German Research Foundation (SFB-TR84TP B08) and the Einstein Foundation Berlin (EP-2017-393).

## Author contributions

C.A.F. conceived the study. G.V., M.V., O.Z-M., L.J.V.G., M.A.M. & C.A.F. designed experiments and analysed the data. G.V., A.R.P., A. Guequén., A. Gianotti, O.Z-M. and C.A.F. performed electrophysiological measurements and data analysis. A.R.P. & L.A. performed CBF determinations and data analysis. G.V. & A. Guequén performed *ex vivo* MCC determinations and data analysis. G.V. performed MCC *in vivo*, BALF, IHQ, histology and data analysis. G.V., M.A.M. & C.A.F. wrote the manuscript, and all authors revised and approved the final version.

---

**SUPLEMENTAL FIGURE 1.**
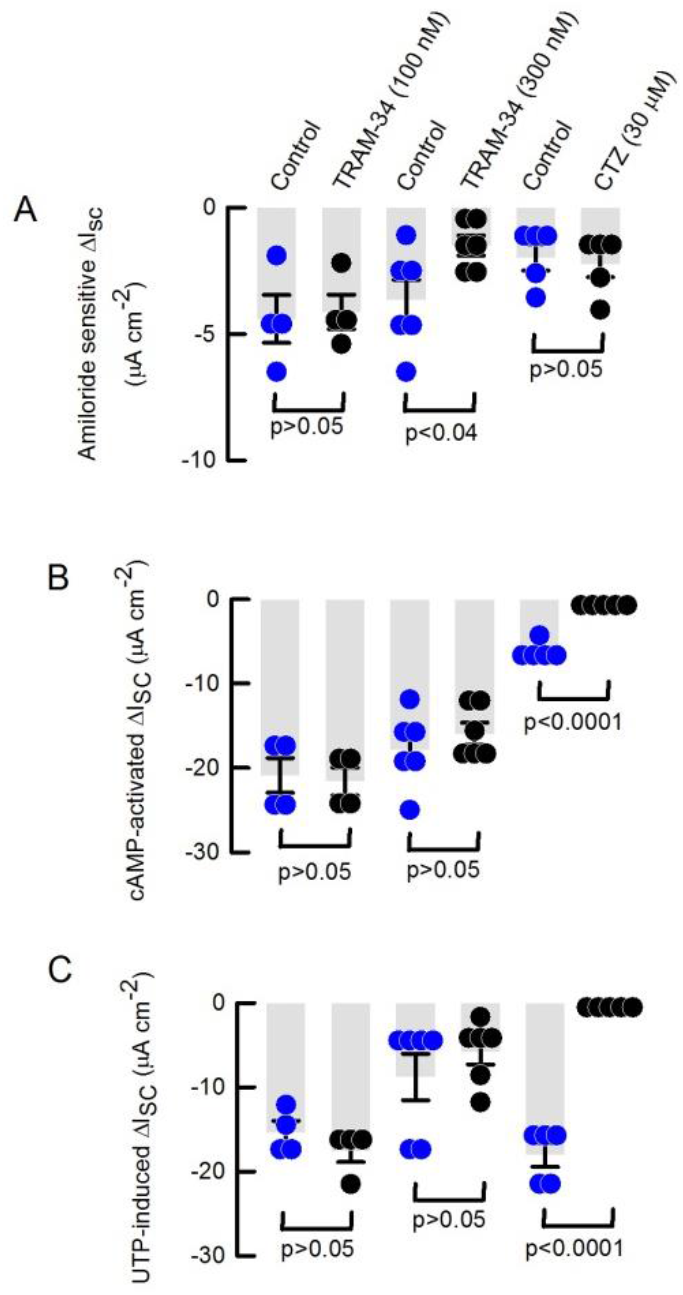
The KCa3.1 inhibitors TRAM-34 and clotrimazol (CTZ), have different effects on short-circuit currents measured in HBECs. (A) summarizes the amiloride-sensitive sodium-absorption. (B) cAMP and (C) Ca^2+^-induced anion secretion was affected by CTZ only. All groups correspond to paired experiments. Calculation of the cAMP-induced I_SC_ in the CTZ and corresponding controls were calculated using the CFTR_inh_172 after cAMP. Analysis performed using Rank sum test.

**SUPLEMENTAL FIGURE 2.**
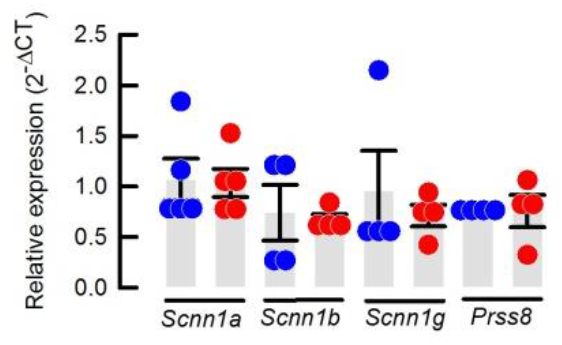
Silencing of *Kcnn4* does not induce changes in the expression of ENaC subunits or prostasin (*Prss8*). (A) summarizes the amiloride-sensitive sodium-absorption. (B) cAMP and (C) Ca^2+^-induced anion secretion was affected by CTZ only. All groups correspond to paired experiments. Calculation of the cAMP-induced I_SC_ in the CTZ and corresponding controls were calculated using the CFTR_inh_172 after cAMP. Analysis performed using a Rank sum test.

